## supplementary figures and tables for "Functional Plasticity in Chromosome-Microtubule Coupling on the Evolutionary Time Scale"

**Supporting figures and tables**

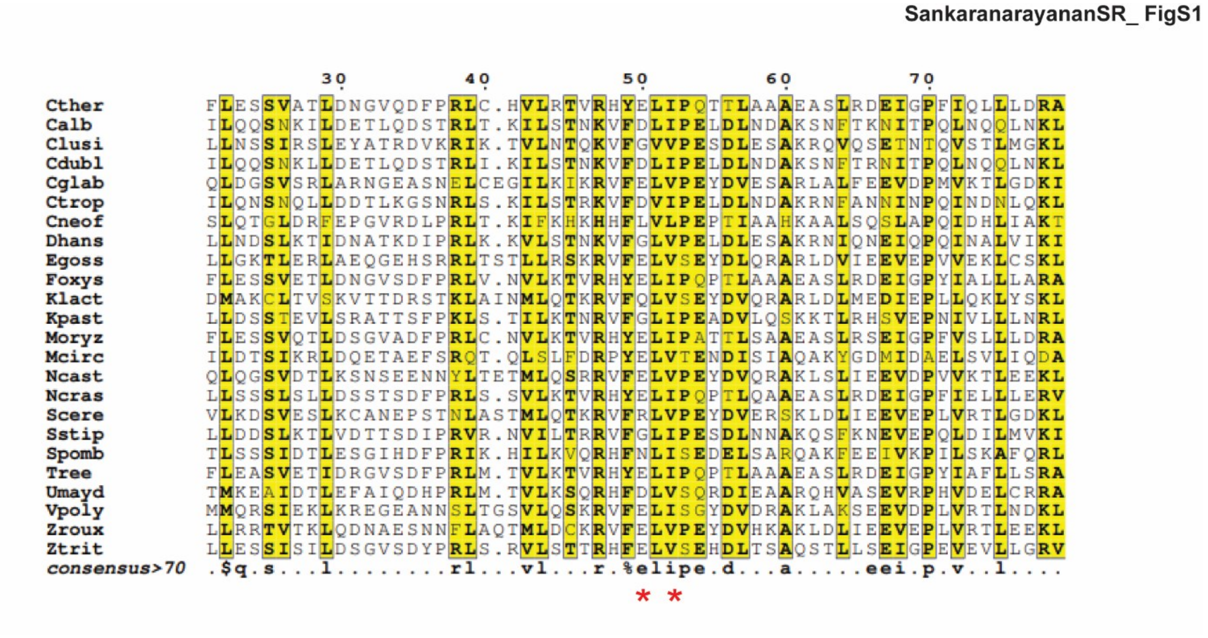

**Figure S1. Multiple sequence alignment showing the conservation of regions in Spc19 predicted to interact with the DSS.** A representative alignment of Spc19 amino acid sequences from species that represent all fungal centromere structures. The residue numbers in the top is based on *Chaetomium thermophilum,* from which we identified residues E50 and I52 to interact with DSS. The position corresponding to these two residues are indicated by a red asterisk in the bottom.

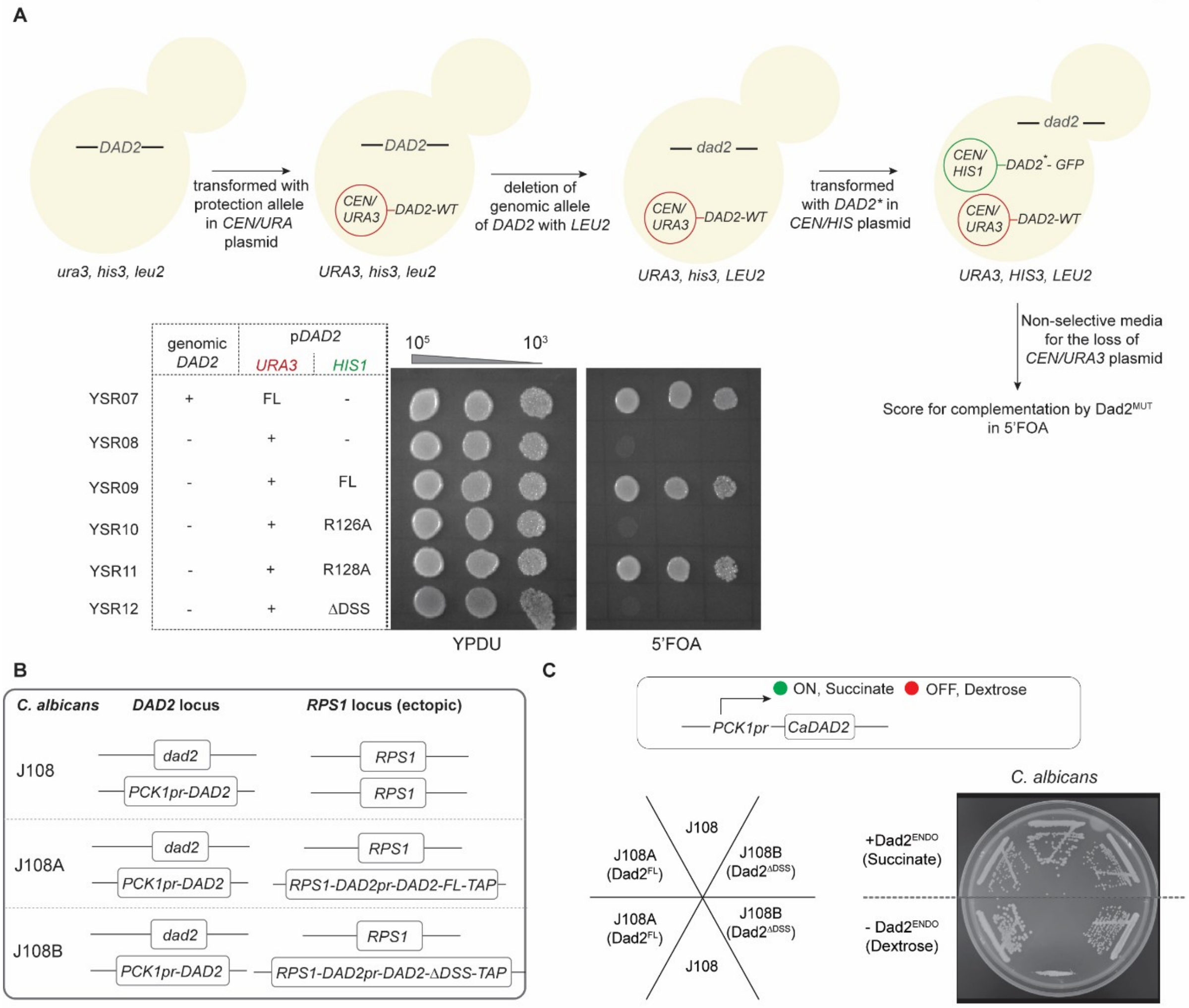

**Figure S2. Essentiality of the DSS is not conserved across point and regional centromeres.** (A) Schematic depicting the construction of strains YSR07 through YSR12 engineered to test for essentiality of the DSS for viability of *S. cerevisiae* at 30°C. The genomic allele of *DAD2* was deleted after protecting the essential Dad2 function with ScDad2-FL from the plasmid pRS316 (*CEN/URA3*). This strain was then transformed with pRS313 (*CEN/HIS3*) containing GFP-tagged full length or the mutant versions of *DAD2* to create strains YSR09 through YSR12. These strains were grown on nonselective media for 12 h, serially diluted ten-fold ($10^5$ to $10^3$), and cells were spotted on YPDU and YPDU+5FOA plates. The plates were incubated at 30°C and imaged after 48 h. (B) The genomic locus of endogenous and ectopic *DAD2* in strains J108, J108A, and J108B engineered to test for the DSS function in *C. albicans* is depicted in a line diagram. (C) Cells from the parent strain J108 (no ectopic CaDad2) along with reintegrant strains J108A (ectopic CaDad2-FL) and J108B (ectopic CaDad2-ΔDSS) were streaked on plates with media permissive or non-permissive for the *PCK1* promoter-driven expression. Plates were imaged post-incubation at 30°C for 48 h. The box above the plate photograph schematically represents the regulation of *PCK1* promoter-driven expression.

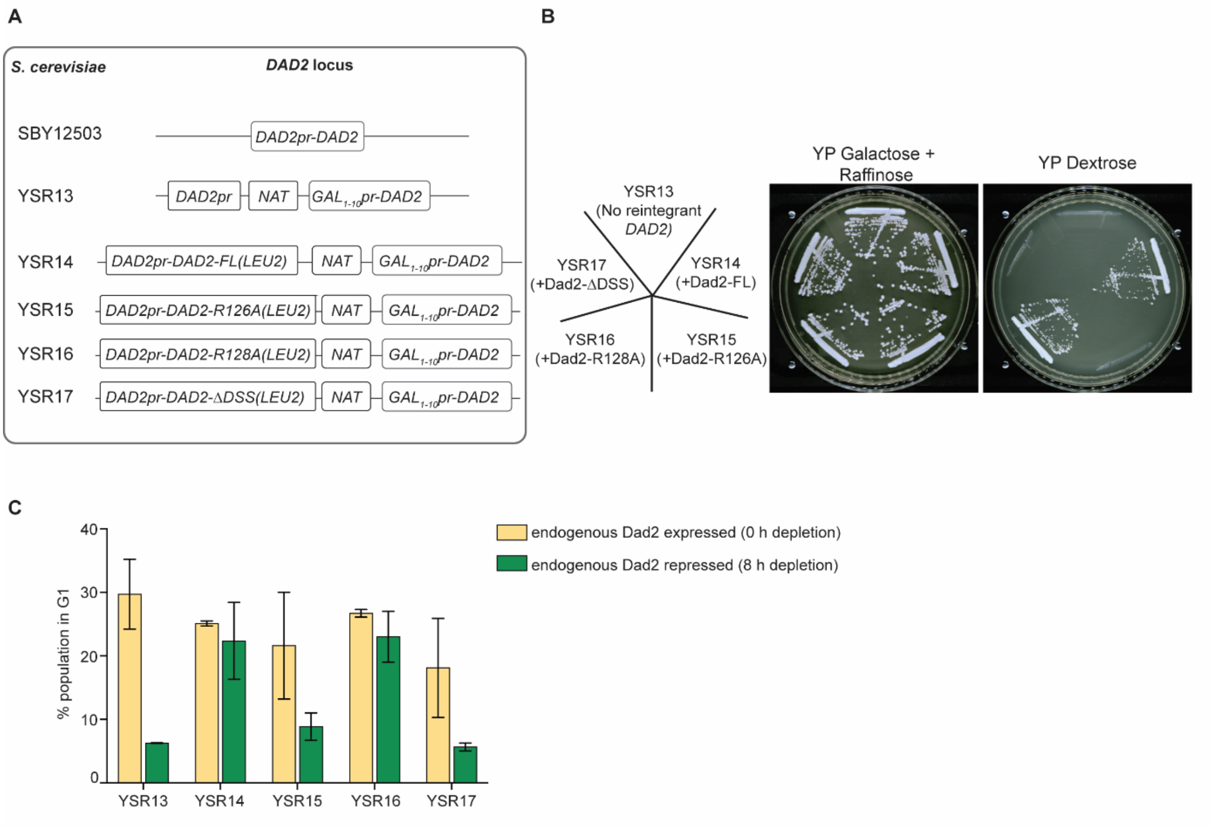

**Figure S3. Mitotic progression in *S. cerevisiae* is dependent on the conserved arginine residue R126.** (A) Line diagrams depict the genomic locus of *DAD2* in the strains SBY12503, YSR13, and the reintegrant strains YSR14 through YSR17 engineered to test for the DSS function in *S. cerevisiae*. (B) Cells from the Dad2 conditional mutant (YSR13) and reintegrant strains expressing indicated versions of Dad2 (YSR14 through YSR17) were streaked on plates with media permissive and non-permissive for expression of the *GAL*$_{1-10}$ promoter that drives the expression of endogenous Dad2. The plates were photographed after incubation at 30°C for 48 h. (C) The proportion of cells in G1 stage in the presence of indicated versions of Dad2 after depleting the endogenous Dad2 for 8h is plotted (N= 30000 events per experiment). Mitotic arrest phenotype in the mutants YSR13, YSR15 and YSR17 is seen to result in lower percentage of cells in G1 as expected.

SankaranarayananSR_ FigS4

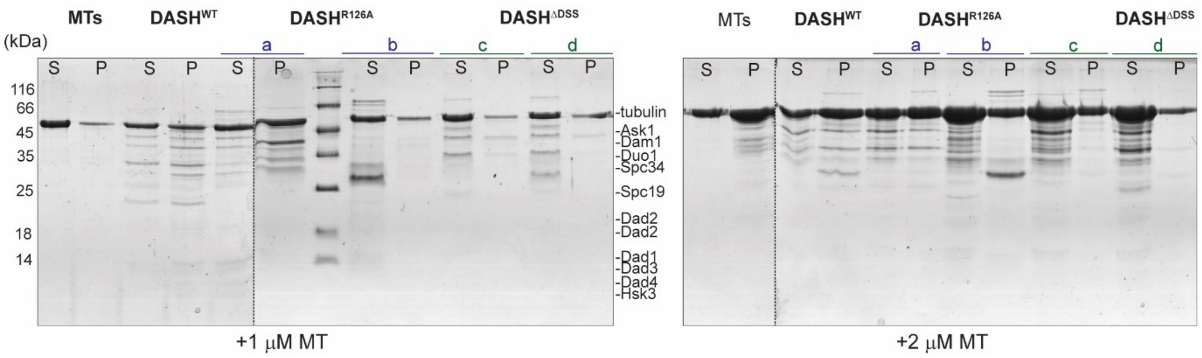

**FigureS4. DSS is essential for Dam1 complex to bind to MTs.** Coomassie stained gels of the cosedimentation assay to test the ability of indicated versions of Dam1 complex with 1μM and 2μM MTs. Lanes marked S and P correspond to supernatant and pellet fractions from the assay. M, molecular weight marker.

SankaranarayananSR_ FigS5

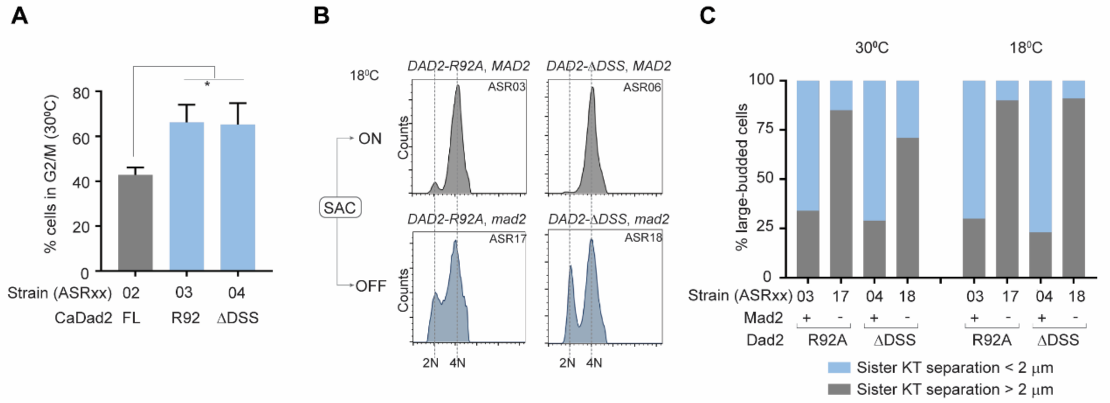

**Figure S5. The conserved DSS is essential for timely cell cycle progression in *C. albicans.*** (A) The bar plot depicts the percentage of cells in the G2/M stage in the indicated strains grown at 30°C (from Figure 3B). The percentage of G2/M cells in each strain was estimated using the cell cycle analysis tool in the software FCS Express 7. Statistical significance was tested by one-way ANOVA (*$p < 0.013$, N=3, 30000 cells per experiment). (B) Histograms depict the distribution of cells with 2N and 4N DNA content (*x*-axis) in the indicated strains after growth at 18°C as analyzed by flow cytometry. ON and OFF respectively indicate the ability or inability of strains to activate the spindle assembly checkpoint (SAC). (C) The percentage of large-budded cells with sister kinetochore separation less than 2 μm (blue) or greater than 2 μm (gray) in the indicated strains upon growth at 30°C and 18°C is plotted. The signs + and – in the *x*-axis indicate the presence or absence of *MAD2* in each strain respectively.

SankaranarayananSR_ FigS6

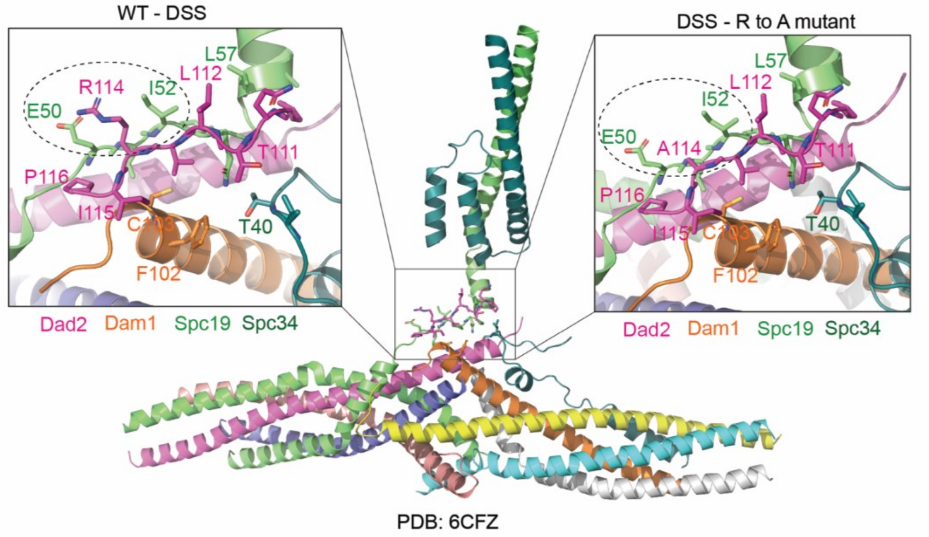

**Figure S6. Neighborhood of the DSS and the conserved arginine residue based on the known structure of the Dam1 complex (PDB 6CFZ).** The location of the DSS in the Dam1 complex monomer is highlighted in the box. The left inset represents the wild-type DSS motif. The residues making steric contacts with the conserved arginine in the DSS (R114 in *Chaetomium thermophilum* Dad2) is shown in dashed circle. The right inset represents the alanine susbsitution of the conserved arginine (A114) and the potential loss of interactions with the residues E50 and I52 of Spc19.

**A**

| Organism | *CEN* structure | CENPA chromatin | CENPA nucleosomes/CEN | kMTs/Chr | Reference |
|---|---|---|---|---|---|
| *S. cerevisiae* | point | 125 bp | 1-2 | 1 | Winey et al.,(1995) |
| *C. albicans* | small regional | 3-5 kb | 4 | 1 | Joglekar et al.,(2008) |
| *S. pombe* | large regional | ~10 kb | 10-15 | 2-3 | Ding et al.,(1993) |
| *G. gallus* | large regional | >30 kb | ND | ~4 | Ribeiro et al.,(2009) |
| *H. sapiens* | large regional | 0.3-5 Mb | ~100 | 12-23 | Wendell et al.,(1993) |
| *C. neoformans* | large regional | 27-64 kb | >10 | >2 | proposed in this study |

**B**

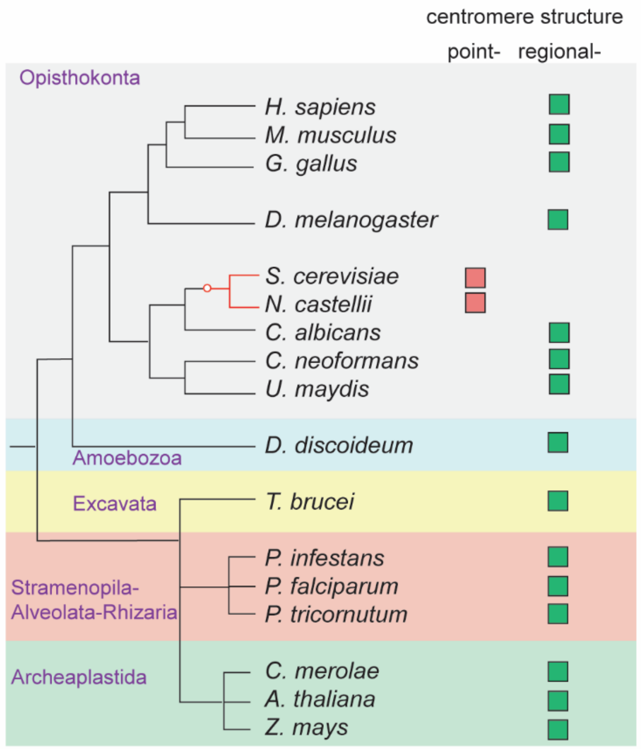

**Figure S7. Regional centromeres can bind multiple kMTs to a chromosome.** (A) A table summarizing experimentally validated centromeric features of common model organisms used to predict the number of CENPA nucleosomes and the number of associated kMTs (red letters) in C. neoformans. (B) The prevalence of regional centromere structure among eukaryotes is illustrated with a cladogram depicting major eukaryotic lineages and the occurrence of point and regional centromere structures in each of them. The cladogram was generated with species representative of each eukaryotic supergroup (adapted from Van Hoofe et al. 2017). The red circle in the cladogram marks the only known origin of point centromeres.

**Table S1. Distribution of point and regional centromere structures across Eukaryota.**

| Taxonomic position | | Species | *CEN* structure | *CEN* size | Ref |
|---|---|---|---|---|---|
| Opisthokonta | Fungi (Ascomycota) | *Saccharomyces cerevisiae* | point | 125 bp | (Fitzgerald-Hayes et al. 1982) |
| | | *Naumovozyma castellii* | point | 125 bp | (Kobayashi et al. 2015) |
| | | *Naumovozyma dairenensis* | point | 125 bp | (Kobayashi et al. 2015) |
| | | *Saccharomyces bayanus* | point | 125 bp | (Gordon et al. 2011) |
| | | *Candida glabrata* | point | 125 bp | (Gordon et al. 2011) |
| | | *Vanderwalozyma polyspora* | point | 125 bp | (Gordon et al. 2011) |
| | | *Zygosaccharomyces rouxii* | point | 125 bp | (Gordon et al. 2011) |
| | | *Lachancea kluveri* | point | 125 bp | (Gordon et al. 2011) |
| | | *Ashbya gossypii* | point | 125 bp | (Gordon et al. 2011) |
| | | *Kluyveromyces lactis* | point | 125 bp | (Gordon et al. 2011) |
| | | *Candida albicans* | regional | 3-5 kb | (Sanyal et al. 2004) |
| | | *Candida dubliniensis* | regional | 3-5 kb | (Padmanabhan et al. 2008) |
| | | *Candida tropicalis* | regional | 10-18 kb | (Chatterjee et al. 2016) |
| | | *Candida parapsilosis* | regional | 1-2.6 kb | (Guin et al. 2020a; Ola et al. 2020) |
| | | *Candida viswanathii* | regional | 11-16 kb | (Guin et al. 2020a) |
| | | *Candida sojae* | regional | 7-26 kb | (Guin et al. 2020a) |
| | | *Candida auris* | regional | 2-3 kb | (Narayanan et al. 2021) |
| | | *Candida lusitaniae* | regional | 4-4.6 kb | (Kapoor et al. 2015) |
| | | *Candida haemulonii* | regional | 2-3 kb | (Narayanan et al. 2021) |
| | | *Candida duobushaamulonii* | regional | 2-3 kb | (Narayanan et al. 2021) |

| | | | | | |
|---|---|---|---|---|---|
| | | *Candida pseudohaemulonii* | regional | 2-3 kb | (Narayanan et al. 2021) |
| | | *Kuraishia capsulata* | regional | 3.5-6 kb | (Marie-Nelly et al. 2014) |
| | | *Ogataea polymorpha* | regional | <10 kb | (Ravin et al. 2013) |
| | | *Blastobotrys adeninivorans* | regional | ~6 kb | (Kunze et al. 2014) |
| | | *Yarrowia lipolytica* | regional | <200 bp | (Fournier et al. 1993) |
| | | *Komgaetaella phaffii* | regional | 3-5 kb | (Coughlan et al. 2016) |
| | | *Schefferosomyces. stipites* | regional | ~10 kb | (Coughlan and Wolfe 2019) |
| | | *Zymoseptoria tritici* | regional | 5-12.5 kb | (Schotanus et al. 2015) |
| | | *Neurospora crassa* | regional | 170-300 kb | (Cambareri et al. 1998) |
| | | *Magnaporthe oryzae* | regional | 57-109 kb | (Yadav et al. 2019) |
| | | *Schizosaccharomyces pombe* | regional | 35-110 kb | (Nakaseko et al. 1986; Fishel et al. 1988) |
| | Fungi (Basidiomycota) | *Cryptococcus neoformans* | regional | 27-64 kb | (Janbon et al. 2014; Yadav et al. 2018) |
| | | *Cryptococcus deuterogattii* | regional | 8-22 kb | (Janbon et al. 2014; Yadav et al. 2018) |
| | | *Malassezia sympodialis* | regional | 3-5.5 kb | (Sankaranarayanan et al. 2020) |
| | | *Malassezia furfur* | regional | 3-5.5 kb | (Sankaranarayanan et al. 2020) |
| | | *Ustilago maydis* | regional | 7-38 kb | (Yadav et al. 2018) |
| | Fungi (Mucoromycota) | *Mucor circinelloides* | mosaic of point and regional | ~1 kb | (Navarro-Mendoza et al. 2019) |
| | Animals | *Drosophila melanogaster* | regional | 200-500 kb | (Sun et al. 1997) |
| | | *Equus asinus* | regional | 54-345 kb | (Nergadze et al. 2018) |

| | | | | | |
|---|---|---|---|---|---|
| | | *Gallus gallus* | regional | >30 kb | (Shang et al. 2010) |
| | | *Mus musculus* | regional | 300-500 kb | (Kipling et al. 1991) |
| | | *Homo sapiens* | regional | 0.3-5 Mb | (Lo et al. 2001) |
| Amoebozoa | Amoeba (Slime mold) | *Dictyostelium discoideum* | regional | 171-361 kb | (Glockner and Heidel 2009) |
| Stramenopila-Alveoloata-Rhizaria (SAR) | Stramenopiles | *Phaeodactylum tricornutum* | regional | 2-5.6 kb | (Diner et al. 2017) |
| | | *Phytopthora sojae* | regional | 211-356 kb | (Fang et al. 2020) |
| | Alveolates | *Plasmodium falciparum* | regional | 2-2.5 kb | (Kelly et al. 2006) |
| | | *Toxoplasma gondii* | regional | 13-20 kb | (Brooks et al. 2011) |
| Archeaplastida | Red algae | *Cyanidioschyzon merolae* | regional | 1-4 kb | (Kanesaki et al. 2015) |
| | Land plants (Angiosperms) | *Oryza sativa* | regional | 420-820 kb | (Yan et al. 2008) |
| | | *Arabidopsis thaliana* | regional | 40-148 kb | (Maluszynska and Heslop-Harrison 1991) |
| | | *Brassica campestris* | regional | 20.2-10 Mb | (Harrison and Heslop-Harrison 1995) |
| | | *Sorghum bicolor* | regional | ND | (Jiang et al. 1996) |
| | | *Triticum aestivum* | regional | ND | (Kishii et al. 2001) |
| | | *Zea mays* | regional | 2-10 Mb | (Ananiev et al. 1998) |
| | | *Solanum tuberosum* | regional | 1-3 Mb | (Gong et al. 2012) |
| Excavata | Kinetoplastid | *Trypanosoma bruceii* | regional | 20-120 kb | (Obado et al. 2007; Echeverry et al. 2012) |

ND, not determined

**Table S2- List of strains used in this study**

| Strain | Parent | Genotype |
|---|---|---|
| ***S. cerevisiae strains*** | | |
| CJY077 | | *MATa Δdad2::KanMX6 ura3-52 lys2-801 ade2-101 trp1Δ63 leu2Δ1::pCJ055(*$DAD2^{TS}$*, LEU2) his3Δ200* [23] |
| YSR01 | CJY077 | *MATa Δdad2::KanMX6 ura3-52 lys2-801 ade2-101 trp1Δ63 leu2Δ1::pCJ055 his3Δ200 pRS313G (GFP, HIS3)* |
| YSR02 | CJY077 | *MATa Δdad2::KanMX6 ura3-52 lys2-801 ade2-101 trp1Δ63 leu2Δ1::pCJ055 his3Δ200 pSR01 (DAD2-FL-GFP, HIS3,CEN6)* |
| YSR03 | CJY077 | *MATa Δdad2::KanMX6 ura3-52 lys2-801 ade2-101 trp1Δ63 leu2Δ1::pCJ055 his3Δ200 pSR02 (DAD2-R126A-GFP, HIS3,CEN6)* |
| YSR04 | CJY077 | *MATa Δdad2::KanMX6 ura3-52 lys2-801 ade2-101 trp1Δ63 leu2Δ1::pCJ055 his3Δ200 pSR03 (DAD2-R128A-GFP, HIS3,CEN6)* |
| YSR05 | CJY077 | *MATa Δdad2::KanMX6 ura3-52 lys2-801 ade2-101 trp1Δ63 leu2Δ1::pCJ055 his3Δ200 pSR04 (DAD2-ΔDSS-GFP, HIS3,CEN6)* |
| BY4741 | | *MATa his3Δ1 leu2Δ0 met15Δ0 ura3Δ0* |
| YSR06 | BY4741 | *MATa his3Δ1 leu2Δ0 met15Δ0 ura3Δ0 SPC42::SPC42-mCherry-KanMX4* |
| YSR07 | YSR06 | *MATa his3Δ1 leu2Δ0 met15Δ0 ura3Δ0 SPC42::SPC42-mCherry-KanMX4 pSR05(Dad2-FL,URA3,CEN6)* |
| YSR08 | YSR07 | *MATa his3Δ1 leu2Δ0 met15Δ0 ura3Δ0 SPC42::SPC42-mCherry-KanMX4 Δdad2::LEU2 pSR05(Dad2-FL,URA3,CEN6)* |
| YSR09 | YSR08 | *MATa his3Δ1 leu2Δ0 met15Δ0 ura3Δ0 SPC42::SPC42-mCherry-KanMX4 Δdad2::LEU2 pSR05(Dad2*$^{-}$*FL,URA3,CEN6) pSR01 (DAD2-FL-GFP, HIS3,CEN6)* |
| YSR10 | YSR08 | *MATa his3Δ1 leu2Δ0 met15Δ0 ura3Δ0 SPC42::SPC42-mCherry-KanMX4 Δdad2::LEU2 pSR05(Dad2- FL,URA3,CEN6) pSR02 (DAD2-R126A-GFP, HIS3,CEN6)* |
| YSR11 | YSR08 | *MATa his3Δ1 leu2Δ0 met15Δ0 ura3Δ0 SPC42::SPC42-mCherry-KanMX4 Δdad2::LEU2 pSR05(Dad2-FL,URA3,CEN6) pSR03(DAD2-R128A-GFP, HIS3,CEN6)* |
| YSR12 | YSR08 | *MATa his3Δ1 leu2Δ0 met15Δ0 ura3Δ0 SPC42::SPC42-mCherry-KanMX4 Δdad2::LEU2 pSR05(Dad2-FL,URA3,CEN6) pSR04 (DAD2-ΔDSS-GFP, HIS3,CEN6)* |

| | | |
|---|---|---|
| SBY12503 | | *MAT a pCUP1-GFP12-LacI12:HIS3 CEN3::33LacO:KanMX SPC110-mCherry:hphMX HSK3-3V5-IAA7:KanMX bar1-1 ade3Δ (UMBRIET N. et al., Nat Commun, 2014; received from S. Biggins, Fred Hutch Cancer Center)* |
| YSR13 | SBY12503 | *MAT a pCUP1-GFP12-LacI12:HIS3 CEN3::33LacO:KanMX SPC110-mCherry:hphMX HSK3-3V5-IAA7:KanMX bar1-1 ade3Δ dad2::$GAL_{1-10}$prDAD2 (NAT)* |
| YSR14 | YSR13 | *MAT a pCUP1-GFP12-LacI12:HIS3 CEN3::33LacO:KanMX SPC110-mCherry:hphMX HSK3-3V5-IAA7:KanMX bar1-1 ade3Δ dad2::$GAL_{1-10}$prDAD2 (NAT) DAD2pr-DAD2-FL (LEU2)* |
| YSR15 | YSR13 | *MAT a pCUP1-GFP12-LacI12:HIS3 CEN3::33LacO:KanMX SPC110-mCherry:hphMX HSK3-3V5-IAA7:KanMX bar1-1 ade3Δ dad2::$GAL_{1-10}$prDAD2 (NAT) DAD2pr-DAD2-R126A (LEU2)* |
| YSR16 | YSR13 | *MAT a pCUP1-GFP12-LacI12:HIS3 CEN3::33LacO:KanMX SPC110-mCherry:hphMX HSK3-3V5-IAA7:KanMX bar1-1 ade3Δ dad2::$GAL_{1-10}$prDAD2 (NAT) DAD2pr-DAD2-R128A (LEU2)* |
| YSR17 | YSR13 | *MAT a pCUP1-GFP12-LacI12:HIS3 CEN3::33LacO:KanMX SPC110-mCherry:hphMX HSK3-3V5-IAA7:KanMX bar1-1 ade3Δ dad2::$GAL_{1-10}$prDAD2 (NAT) DAD2pr-DAD2-ΔDSS (LEU2)* |
| ***C.albicans strains*** | | |
| SN148 | | *Δura3::imm434/Δura3::imm434,Δhis1::hisG/Δhis1::hisG Δarg4::hisG/Δarg4::hisG, Δleu2::hisG/Δleu2::hisG* (Noble and Johnson, 2005) |
| J108 | | *Δura3::imm434/Δura3::imm434,Δhis1::hisG/Δhis1::hisG Δarg4::hisG/Δarg4::hisG dad2::HIS1/PCK1pr-DAD2 (URA3)* (Thakur and Sanyal, 2011) |
| J108A | J108 | *Δura3::imm434/Δura3::imm434,Δhis1::hisG/Δhis1::hisG Δarg4::hisG/Δarg4::hisG dad2::HIS1/PCK1pr-DAD2 (URA3) RPS1/RPS1:DAD2pr-$DAD2^{FL}$-TAP(NAT)* |
| J108B | J108 | *Δura3::imm434/Δura3::imm434,Δhis1::hisG/Δhis1::hisG Δarg4::hisG/Δarg4::hisG dad2::HIS1/PCK1pr-DAD2 (URA3) RPS1/RPS1:DAD2pr-$DAD2^{\Delta DSS}$-TAP(NAT)* |
| ASR01 | SN148 | *Δura3::imm434/Δura3::imm434,Δhis1::hisG/Δhis1::hisG Δarg4::hisG/Δarg4::hisG, Δleu2::hisG/Δleu2::hisG dad2::HIS1/DAD2* |
| ASR02 (CaDad2-FL) | ASR01 | *Δura3::imm434/Δura3::imm434,Δhis1::hisG/Δhis1::hisG Δarg4::hisG/Δarg4::hisG, Δleu2::hisG/Δleu2::hisG dad2::HIS1/DAD2-FL-GFP(URA3)* |

| | | |
|---|---|---|
| ASR03 (CaDad2-R92A) | ASR01 | *Δura3::imm434/Δura3::imm434,Δhis1::hisG/Δhis1::hisG Δarg4::hisG/Δarg4::hisG, Δleu2::hisG/Δleu2::hisG dad2::HIS1/DAD2-R92A-GFP(URA3)* |
| ASR04 (CaDad2-ΔDSS) | ASR01 | *Δura3::imm434/Δura3::imm434,Δhis1::hisG/Δhis1::hisG Δarg4::hisG/Δarg4::hisG, Δleu2::hisG/Δleu2::hisG dad2::HIS1/DAD2-ΔDSS-GFP(URA3)* |
| ASR07 (CaDad2-FL,CENP-A-TAP) | ASR02 | *Δura3::imm434/Δura3::imm434,Δhis1::hisG/Δhis1::hisG Δarg4::hisG/Δarg4::hisG, Δleu2::hisG/Δleu2::hisG dad2::HIS1/DAD2-FL-GFP(URA3) CSE4/CSE4-TAP (LEU2)* |
| ASR08 (CaDad2-R92A, CENP-A-TAP) | ASR03 | *Δura3::imm434/Δura3::imm434,Δhis1::hisG/Δhis1::hisG Δarg4::hisG/Δarg4::hisG, Δleu2::hisG/Δleu2::hisG dad2::HIS1/DAD2-R92A-GFP(URA3)* |
| ASR09 (CaDad2-ΔDSS, CENP-A-TAP) | ASR04 | *Δura3::imm434/Δura3::imm434,Δhis1::hisG/Δhis1::hisG Δarg4::hisG/Δarg4::hisG, Δleu2::hisG/Δleu2::hisG dad2::HIS1/DAD2-ΔDSS-GFP(URA3) CSE4/CSE4-TAP (LEU2)* |
| ASR12 (CaDad2-FL, Tub4-mCherry) | ASR02 | *Δura3::imm434/Δura3::imm434,Δhis1::hisG/Δhis1::hisG Δarg4::hisG/Δarg4::hisG, Δleu2::hisG/Δleu2::hisG dad2::HIS1/DAD2-FL-GFP(URA3) TUB4/TUB4-mCherry(NAT)* |
| ASR13 (CaDad2-R92A, Tub4-mCherry) | ASR03 | *Δura3::imm434/Δura3::imm434,Δhis1::hisG/Δhis1::hisG Δarg4::hisG/Δarg4::hisG, Δleu2::hisG/Δleu2::hisG dad2::HIS1/DAD2-R92A-GFP(URA3) TUB4/TUB4-mCherry(NAT)* |
| ASR14 (CaDad2-ΔDSS, Tub4-mCherry) | ASR04 | *Δura3::imm434/Δura3::imm434,Δhis1::hisG/Δhis1::hisG Δarg4::hisG/Δarg4::hisG, Δleu2::hisG/Δleu2::hisG dad2::HIS1/DAD2-ΔDSS-GFP(URA3) TUB4/TUB4-mCherry(NAT)* |
| ASR03M | ASR03 | *Δura3::imm434/Δura3::imm434,Δhis1::hisG/Δhis1::hisG Δarg4::hisG/Δarg4::hisG, Δleu2::hisG/Δleu2::hisG dad2::HIS1/DAD2-R92A-GFP(URA3) mad2::LEU2/MAD2* |
| ASR17 (CaDad2-R92A, *mad2*) | ASR03M | *Δura3::imm434/Δura3::imm434,Δhis1::hisG/Δhis1::hisG Δarg4::hisG/Δarg4::hisG, Δleu2::hisG/Δleu2::hisG dad2::HIS1/DAD2-R92A-GFP(URA3) mad2::LEU2/mad2::ARG4* |
| ASR04M | ASR04 | *Δura3::imm434/Δura3::imm434,Δhis1::hisG/Δhis1::hisG Δarg4::hisG/Δarg4::hisG, Δleu2::hisG/Δleu2::hisG dad2::HIS1/DAD2-ΔDSS-GFP(URA3) mad2::LEU2/MAD2* |
| ASR18 (CaDad2-ΔDSS, *mad2*) | ASR04M | *Δura3::imm434/Δura3::imm434,Δhis1::hisG/Δhis1::hisG Δarg4::hisG/Δarg4::hisG, Δleu2::hisG/Δleu2::hisG dad2::HIS1/DAD2-ΔDSS-GFP(URA3) mad2::LEU2/mad2::ARG4* |

| | | |
|---|---|---|
| ASR19 (CaDad2-FL, Ndc80-mCherry) | ASR02 | *Δura3::imm434/Δura3::imm434,Δhis1::hisG/Δhis1::hisG Δarg4::hisG/Δarg4::hisG, Δleu2::hisG/Δleu2::hisG dad2::HIS1/DAD2-FL-GFP(URA3) NDC80/NDC80-mCherry-ARG4* |
| ASR20 (CaDad2-R92A, Ndc80-mCherry) | ASR03 | *Δura3::imm434/Δura3::imm434,Δhis1::hisG/Δhis1::hisG Δarg4::hisG/Δarg4::hisG, Δleu2::hisG/Δleu2::hisG dad2::HIS1/DAD2-R92A-GFP(URA3) NDC80/NDC80-mCherry-ARG4* |
| ASR21 (CaDad2-ΔDSS, Ndc80-mCherry) | ASR04 | *Δura3::imm434/Δura3::imm434,Δhis1::hisG/Δhis1::hisG Δarg4::hisG/Δarg4::hisG, Δleu2::hisG/Δleu2::hisG dad2::HIS1/DAD2-ΔDSS-GFP(URA3) NDC80/NDC80-mCherry-ARG4* |
| ***C. neoformans strains*** | | |
| CNVY108 | | *α H99::GFP-H4-NAT (pVY3) (Kozubowski et al., mBio, 2013)* |
| SHR741 | | *α H99::GFP-H4-NAT, mad2::NEO (Sridhar et al., Nat. Commun., 2021)* |
| CNSD169 | CNVY108 | *α H99::GFP-H4-NAT DAD2::DAD2p-DAD2-mCherry-NEO* |
| CNSD170 | CNVY108 | *α H99::GFP-H4-NAT DAD2::DAD2p-DAD2-R102A-mCherry-NEO* |
| CNSD171 | CNVY108 | *α H99::GFP-H4-NAT DAD2::DAD2p-DAD2-ΔDSS-mCherry-NEO* |
| CNVY104 | | *α H99::GFP-DAD1-NAT (pVY2) (Kozubowski et al., mBio, 2013)* |
| CNSD198 | CNSD169 | *α H99::GFP-DAD1-NAT (pVY2) DAD2::DAD2p-DAD2 -mCherry-NEO* |
| CNSD199 | CNSD170 | *α H99::GFP-DAD1-NAT (pVY2) DAD2::DAD2p-DAD2-R102A-mCherry-NEO* |
| CNSD200 | CNSD171 | *α H99::GFP-DAD1-NAT (pVY2) DAD2::DAD2p-DAD2-ΔDSS-mCherry-NEO* |
| ***S. pombe strains*** | | |
| SP22 | | *h- ade6-M210 leu1 lys1 ura4 cen2(D107)::KanR -ura4-lacO his7+::lacI-GFP (received from A. Marston, University of Edinburgh)* |
| SP1619 | SP22 | *h- ade6-M210 leu1 lys1 ura4 cen2(D107)::KanR -ura4-lacO his7+::lacI-GFP dad2 ::HPH (received from A. Marston, University of Edinburgh)* |
| PSR01 | SP1619 | *h- ade6-M210 leu1 lys1 ura4 cen2(D107)::KanR -ura4-lacO his7+::lacI-GFP dad2 ::HPH DAD2pr::DAD2pr-DAD2-FL-FLAG-NAT* |

| | | |
|---|---|---|
| PSR02 | SP1619 | *h- ade6-M210 leu1 lys1 ura4 cen2(D107)::KanR -ura4-lacO his7+::lacI-GFP dad2 ::HPH DAD2pr::DAD2pr-DAD2-ΔDSS-FLAG-NAT* |

**Table S3- List of plasmids used in this study**

| **Plasmid** | **Parent** | ***Description*** |
|---|---|---|
| pRS313 | | *CEN6/ARS/HIS3 plasmid* |
| pRS313G | pRS313 | *GFP along with CYC1 terminator cloned into pRS313 in BamHI-ClaI sites* |
| pSR01 | pRS313G | *ScDAD2-FL cloned in pRS313G in SacII-BamHI sites* |
| pSR02 | pRS313G | *ScDAD2-R126A cloned in pRS313G in SacII-BamHI sites* |
| pSR03 | pRS313G | *ScDAD2-R128A cloned in pRS313G in SacII-BamHI sites* |
| pSR04 | pRS313G | *ScDAD2-ΔDSS cloned in pRS313G in SacII-BamHI sites* |
| pRS316 | | *CEN6/ARS/URA3 plasmid* |
| pSR05 | pRS316 | *ScDAD2-FL with native promoter and terminator cloned in pRS316 in SacII-SacI sites* |
| pYM-N25 | | *Plasmid used to amplify cassette to place DAD2 under $GAL_{1-10}$ promoter with NAT marker* |
| pUG73 | | *Plasmid to amplify DAD2 deletion cassette with LEU2 marker, also used to generate reintegration plasmids* |
| pSR06 | pUG73 | *ScDAD2-FL cloned in SacII-SacI sites* |
| pSR07 | pUG73 | *ScDAD2-R126A cloned in SacII-SacI sites* |
| pSR08 | pUG73 | *ScDAD2-R128A cloned in SacII-SacI sites* |
| pSR09 | pUG73 | *ScDAD2-ΔDSS cloned in SacII-SacI sites* |
| pBS-RN | pBS-NAT | *CaRPS1 cloned in pBS NAT plasmid in NotI site* |
| pRN-Dad2$^{FL}$ | pBS-RN | *CaDAD2pr-CaDAD2-FL-TAP cloned in SalI site* |
| pRN-Dad2$^{\Delta DSS}$ | pBS-RN | *CaDAD2pr-CaDAD2-*ΔDSS *-TAP cloned in SalI site* |
| pRN-92 | pBS-RN | *CaDAD2pr-CaDAD2-R92A-TAP cloned in SalI site* |
| pCse4TAPLeu | | *CSE4-TAP with LEU2 marker (Varshney and Sanyal, 2019)* |
| pTub4-mCherryNAT | pDam1-mCherryNAT | *TUB4 ORF cloned in pDam1-mCherryNAT in SacII-SpeI sites* |
| pMad2-2 | | *MAD2 deletion cassette with LEU2 (Thakur and Sanyal, 2011)* |
| pMad2-3 | | *MAD2 deletion cassette with ARG4 (Thakur and Sanyal, 2011)* |
| pSR10 | pBS-GFPUra | *3'UTR of CaDAD2 cloned in pBS-GFPUra in XhoI-KpnI sites* |
| pSR11 (CaDad2-FL) | pSR10 | *CaDAD2-FL cloned in pSR10 in SacII-SpeI sites* |
| pSR12 (CaDad2-R'A) | pSR10 | *CaDAD2-R92A cloned in pSR10 in SacII-SpeI sites* |

| | | |
|---|---|---|
| pSR15 (CaDad2-ΔDSS) | pSR10 | *CaDAD2-*ΔDSS *cloned in pSR10 in SacII-SpeI sites* |
| pLK25 | | *Amplification of mCherry-Neomycin for C. neoformans* |
| PC4-Dad1H | | *Polycistronic vector for the expression and purification of Dam1complex from E. coli. (Miranda et al., 2005, procured from addgene)* |
| PC4-126A | PC4-Dad1H | *Dad2 ORF replaced to express Dad2-R126A, to purify the mutant Dam1 complex* |
| PC4-ΔDSS | PC4-Dad1H | *Dad2 ORF replaced to express Dad2-ΔDSS, to purify the mutant Dam1 complex* |
| pVB128 | | *Yeast integrative plasmid for 3xFLAG tagging at the C terminus using NAT marker (received from V. Borde lab, Institut Curie)* |
| pSp-FL | pVB128 | *SpDad2pr-Dad2-FL cloned in frame with 3xFLAG for reintegration into the Dad2 promoter sequence by NheI digestion* |
| pSp-ΔDSS | pVB128 | *SpDad2pr-Dad2-*ΔDSS *cloned in frame with 3xFLAG for reintegration into the Dad2 promoter sequence by NheI digestion* |

**Table S4- List of primers used in this study**

| | | |
|---|---|---|
| ***S. cerevisiae*** | | |
| ScGFP-F | CATGGATCCATGGTGAGCAAGGGCG | Cloning GFP into pRS313 |
| ScGFP-R | GCTGATCGATGCAAATTAAAGCCTTCGAGC | |
| SR282 | TCCCGCGGCATTGCGGCAGGTAAAATATC | Amplification of $Dad2^{FL/MUT}$ as SacII/BamHI fragment to clone into pRS313G |
| ScDad2R | GAGGGATCCTTCGTTACCATCTACCCTAATTCTG | |
| Sc126A-R | GAGGGATCCTTCGTTACCATCTACCCTAATTGCGACCAT TGTTTCC | |
| Sc128A-R | GAGGGATCCTTCGTTACCATCTACGGCAATTCTGACCAT TGTTTCC | |
| ScΔDSS-R | GAGGGATCCTTCGTTACCATCCAAGGGTACCAGATCTT C | |
| ScDad2Pr F | TGACCGCGGGTACAATGGTCCTAACTTAATG | Cloning ScDad2-FL into pRS316 |
| ScDad2R | ATCGAGCTCCCAACACTGTAGAATACTAATATC | |
| ScDad2delF | GAATATATCTAAAAAACTATTGAATAGGTTTGAAAAAC TCATAATTCAGACAGTTATTGCTGTGAAGATCCCAGCA AAGG | Deletion of genomic allele of DAD2 with LEU2 marker |
| ScDad2delR | ATGTATATGCTATTATCGCAACTGCCTTCTTCCGATTTA TATAAGATCCTTTTCTTTCTGTGCAGGCTAACCGGAACC TGT | |
| ScDad2F | GGTTAGAGGGCGACAATAC | Confirmatory primers |
| Leu2R | CACCAGTGTTCAACTCAACAAG | |
| GalDad2FP | CTAAAAAACTATTGAATAGGTTTGAAAAACTCATAATT CAGACAGTTATTCGACATGGAGGCCCAGAATAC | Cassette to place DAD2 under GAL promoter with NAT marker |
| GalDad2RP | GACTGAAGTTCTTTTCGCTTTATAGCAATTTGTTCATCT ATTGAATCCATTTTGTACAATTCATCCATACCATGG | |
| Dad2pr-SacII-F | TGACCGCGGGTACAATGGTCCTAACTTAATG | Overlap PCR primers to amplify $DAD2^{FL/MUT}$ for cloning into pUG73 for reintegration |
| Dad2pr-SacI-R | ATCGAGCTCCCAACACTGTAGAATACTAATATC | |
| ScTer FP | GATGGTAACGAATGAACAGAAAG | |
| Sc126A-R | CTTTCTGTTCATTCGTTACCATCTACCCTAATTGCGACC ATTGTTTCC | |
| Sc128A-R | CTTTCTGTTCATTCGTTACCATCTACGGCAATTCTGACC ATTGTTTCC | |
| ScΔDSS-R | CTTTCTGTTCATTCGTTACCCAAGGGTACCAGATCTTC | |
| ***C. albicans*** | | |
| RP10F | ATAAGAATGCGGCCGCTAGATCCAACTCAAGTAC AACATGGC | amplification and cloning of RPS1 locus into pBS-NAT |
| RP10R | ATAAGAATGCGGCCGCGGATCCCCCAGATCATTA | |

| | | |
|---|---|---|
| | TCC | |
| AD02 | ACGCGTCGACATTTTGACTAGT TCTCAAATGGTTC | Overlap PCR primers to amplify CaDAD2$^{FL/MUT}$ and clone into pBS-RP10-NAT |
| AD03 | CCATCGATCGATATCAAGCTTCAGGTTG | |
| Dad2delR | GTCTTGATTTTCTTCATTTGATTGTCCGAAAGCCTCTTG TTATTATC | |
| Dad2delF | GATAATAACAAAGAGGCTTTCGGACAATCAAATGAAGA AAATCAAGAC | |
| R92A FP | GAACCATTAGTAGCAGTGCGTGTTGGACAATCA | |
| R92A RP | TGATTGTCCAACACGCACTGCTACTAATGGTTC | |
| Dad2DS-F | GGCCTCGAGGTGTACATATAATAACTCTAAATTCTGGC | Amplification of Dad2 3'UTR homology |
| Dad2DS-R | ACCGGTACCGAATTTTGTCAACCAAGAAATAGAC | |
| Dad2FP | TACCCGCGGATGCTGAAAACAAATACTGCTATATACC | Amplification of DAD2 ORF tagging with GFP |
| Dad2GFP-RP | CGACTAGTTTCCGTGGATTCTTCAACTTC | |
| Dad2cFP | TCAATACCCACCACAAAACC | Confirmatory primer |
| ***C. neoformans*** | | |
| SD118 | TGCATGCATTCTCGTCAAAATAGGCTGC | Overlap PCR primers to amplify CnDAD2$^{FL/R102A}$ |
| SD119 | TCGCCGCCGTATGCTAAGGCCACGAGACAAGGGAGAG G | |
| SD120 | CCTCTCCCTTGTCTCGTGGCCTTAGCATACGGCGGCGAA | |
| VYP152 | CTCGCCCTTGCTCACCATTTGTTGTTTTGTTTTATCAGAT GCG | |
| VYP153 | CTGATAAAACAAAACAACAAATGGTGAGCAAGGGCGA G | Amplification of mCherry-NEO from pLK25 |
| VYP154 | CTATTGGTCGTCATCAGCAGGCCAAGCTTGGTACCGAG CTC | |
| VYP155 | GAGCTCGGTACCAAGCTTGGCCTGCTGATGACGACCAA TAG | amplification CnDAD2 3'UTR homology |
| VYP156 | CCCAAGCTTCTCCATATCGTGTCTCAATTTCATCTC | |
| VYP157 | CTTGTACAGCTCGTCCATGC | Confirmatory primer |
| ***S. pombe*** | | |
| VB128 FP | CACCACCATCATCATCACGG | Amplification of the vector pVB128 |
| VB128 RP | CACCACCATCATCATCACGG | |
| SpFP | CGGCCAGTGAATTGTAATACGACTCGGAACGTTGACAA TCTTGTTG | Sp-FP along with each of the reverse primer to amplify |
| Sp-FL-RP | ccgtgatgatgatggtggtgTACCTCTTCAACATCGCCTTGC | |

| | | |
|---|---|---|
| Sp-ΔDSS-RP | ccgtgatgatgatggtggtgTACCTCTTCAACATCGCCTTGCTCCGTAGCAGATGCGTTGGTATTTGAAGTATGTTGGGATGCTATCTGTATTGAC | SpDad2-FL and SpDad2-ΔDSS |
| Sp-cFP | GGTATTCTTTAACGACCCGTTG | Primers to confirm targeted integration |
| FLAG-cRP | GAGGCAAGCTAAACAgatcTC | |

**Table S5- (Attached as an excel sheet)**
